# Adaptive IRE1 Signaling Elicits T Cell Metabolic Remodeling and Tumor Control

**DOI:** 10.1101/2023.11.16.567431

**Authors:** Brian P. Riesenberg, Evelyn J. Gandy, Andrew S. Kennedy, Elizabeth G. Hunt, Katie E. Hurst, Elise L. Sedlacek, Peng Gao, Michael J. Emanuele, Jennfier Modliszewski, Jarred M. Green, Justin J. Milner, R. Luke Wiseman, Jessica E. Thaxton

## Abstract

The efficacy of cancer immunotherapies is limited by the metabolic instability of the tumor microenvironment (TME) that disables T cell antitumor immunity. Metabolic imbalances within the TME are sensed and responded to by stress sensors of the endoplasmic reticulum (ER) unfolded protein response (UPR). The UPR comprises three integrated signaling pathways harboring both adaptive and deleterious phases based on the extent and duration of cell stress. Here, we elucidate the differential contributions of adaptive and deleterious signaling downstream of the UPR IRE1 pathway in T cell-regulated tumor control. T cells in murine and patient cancers experience persistent ER stress, leading to hyperactive IRE1 signaling that limits tumor control. However, amplifying the adaptive arm of the IRE1 UPR serves to eliminate mitochondrial toxicity and protect T cells from chronic ER stress, yielding robust tumor engraftment and long-term tumor immunity. Our findings establish the UPR’s essential protective role in antitumor immunity.

## Introduction

T cell-based cellular immunotherapies such as adoptive cellular therapy and chimeric antigen receptor therapy have shown remarkable clinical success in extending the survival of patients with hematological malignancies^1, 2^. Similar efficacies have not been observed in patients with solid cancer due in part to the highly suppressive nature of the tumor microenvironment (TME)^3^. Thus, a pressing need is to identify T cell-intrinsic mechanisms that disarm antitumor function in the solid TME. Recent pan-cancer T cell atlas data identified that the cell response to stress is a ubiquitous and leading detractor of T cell therapeutic efficacy in tumors^4^. Solid tumors present an onslaught of metabolic stressors reported to impede T cell function in solid cancers including persistent antigen encounter, nutrient deprivation, and hypoxia^5^. Therefore, the intersection between TME stress and the T cell-intrinsic pathways that dynamically control the cellular response to stress is an exciting avenue to improve the efficacy of tumor immunotherapies.

Endoplasmic reticulum (ER) stress was previously shown to be a critical determinant in dictating tumor immunotherapy efficacy^6–8^. Cells sense and respond to ER stress through the activity of the unfolded protein response (UPR) – a stress-signaling pathway comprising of three integrated pathways regulated downstream of the ER membrane proteins inositol-requiring enzyme 1 (IRE1), protein kinase R-like ER kinase (PERK), and activating transcription factor 6 (ATF6). Metabolic perturbations in the TME enforce aberrant protein folding, leading to ER stress and subsequent UPR activation^9^. In response to ER stress, UPR pathways promote the adaptive remodeling of ER proteostasis and cell metabolism to alleviate ER stress and adapt cellular homeostasis^10^. However, in response to chronic, unresolvable ER stress, these pathways promote deleterious apoptotic and pro-inflammatory signaling linked to the pathogenesis of etiologically diverse human diseases^11^.

We and others established that antitumor immunity is subject to hyperactive and persistent UPR activation, primarily through the IRE1 pathway, which dramatically impedes T cell function in multiple solid tumor types^6, 8, 12^. IRE1 regulates the most evolutionarily conserved arm of the UPR and primarily functions through the regulated splicing of mRNA encoding the transcription factor X box binding protein 1 (XBP1s). In ovarian cancers, persistently defective N-linked protein glycosylation in CD4^+^ T cells promotes hyperactive IRE1-XBP1s activity that limits T cell tumor control through dysregulation of metabolism^7^. Similarly, in CD8^+^ tumor infiltrating lymphocytes, cholesterol in the TME prompts chronic IRE1-XBP1s signaling that enforces T cell exhaustion, limiting antitumor immunity^8^. While deletion of IRE1 improves tumor control^13^, it is poorly understood whether the simultaneous loss of the adaptive and protective IRE1 UPR impairs the potential of T cell antitumor potency.

Use of arm-selective UPR activator compounds that act in specific and transient manners has enabled elegant characterization of the adaptive and protective effects of multiple arms of the UPR^14^. Moreover, using transient activators, enhancing adaptive facets of the UPR has served to protect cells from ER stress in disease states that normally induce unresolved ER stress and hyperactive UPR signaling. For example, selective pharmacologic IRE1-XBP1s activation afforded by the compound IXA4 alleviated ER proteostasis imbalances and mitochondrial toxicity associated with the expression of amyloid precursor protein (APP) variants^15^. Further, in obese insulin-resistant mice, systemic treatment with IXA4 was shown to improve global glucose homeostasis and correct metabolic dysfunction in multiple tissues, including steatosis in the liver and insulin regulation in the pancreas^16^. Given that T cells infiltrating solid tumors experience profound mitochondrial toxicity and dysmorphia coupled with severe ER stress-induced metabolic dysfunction^6, 17^, the abovementioned data raise the possibility that enhancing adaptive and protective traits via transient activation of the IRE1 arm of the UPR could protect tumor-infiltrating T cells from ER stress and subsequent dysfunction.

Herein, we pursued the hypothesis that ablation of ER stress sensor IRE1 limits the potential of T cell tumor control by negating the protective arm of the IRE1 UPR. First, by exploiting IRE1^−/−^ CD8^+^ TILs, we established a system to monitor IRE1-linked ER stress using a fluorescent probe that measures proteasome activity. Employing the probe to assess CD8^+^ TILs across multiple tumor models, we identified that highly dysfunctional CD8^+^ TILs undergo chronic ER stress leading to hyperactive IRE1 signaling that limits tumor control. Thus, we observed that IRE1 ablation relieved hyperactive IRE1 signaling, promoting T cells that provided superior tumor control relative to WT CD8^+^ T cells. However, we observed that IRE1^−/−^ T cells had limited engraftment in tumors, suggesting that protective aspects of enhanced survival attributed to the protective phase of the UPR were limited upon IRE1 ablation. The data prompted us to study the contribution of the adaptive and protective phase of the IRE1 UPR to T cell antitumor immunity. CD8^+^ T cells treated with the transient IRE1-XBP1s activator IXA4 underwent robust enrichment of mitochondria complemented by metabolic remodeling that supported amplified engraftment and long-lived tumor immunity. Together, our data establish a novel paradigm for regulating the UPR to cope with ER stress in tumors.

## Results

### Proteasome activity identifies IRE1-regulated ER Stress in CD8^+^ TILs

Previously, we found that T cells in MCA-205 fibrosarcomas were under chronic ER stress evidenced by hyperactive signaling within the PERK axis that impaired T cell-mediated tumor control^6, 12^. We reasoned that other stress sensors could similarly impair T cell function in this tumor model. To identify signaling pathways induced by other ER stress sensors, we performed RNA-sequencing on CD8^+^ T cells isolated from spleens or tumors of MCA-205 fibrosarcoma-bearing mice. Using gene set enrichment analysis (GSEA), we confirmed CD8^+^ T cells from tumors underwent ER stress as they had a significant upregulation of genes involved in pathways associated with the Regulation of Response to ER Stress and the Response to Topologically Incorrect Proteins (Figure 1a). These data substantiated a damaged protein burden in T cells within tumors relative to autologous-matched splenocytes. A deeper analysis of this data found that three of the most significantly enriched genes in the tumor were directly related to the IRE1 pathway, *Hspa5*, *Xbp1*, *Ern1* (the gene encoding IRE1), indicating that IRE1 signaling is impacted in CD8^+^ TILs (Figure 1b).

**Figure 1.**
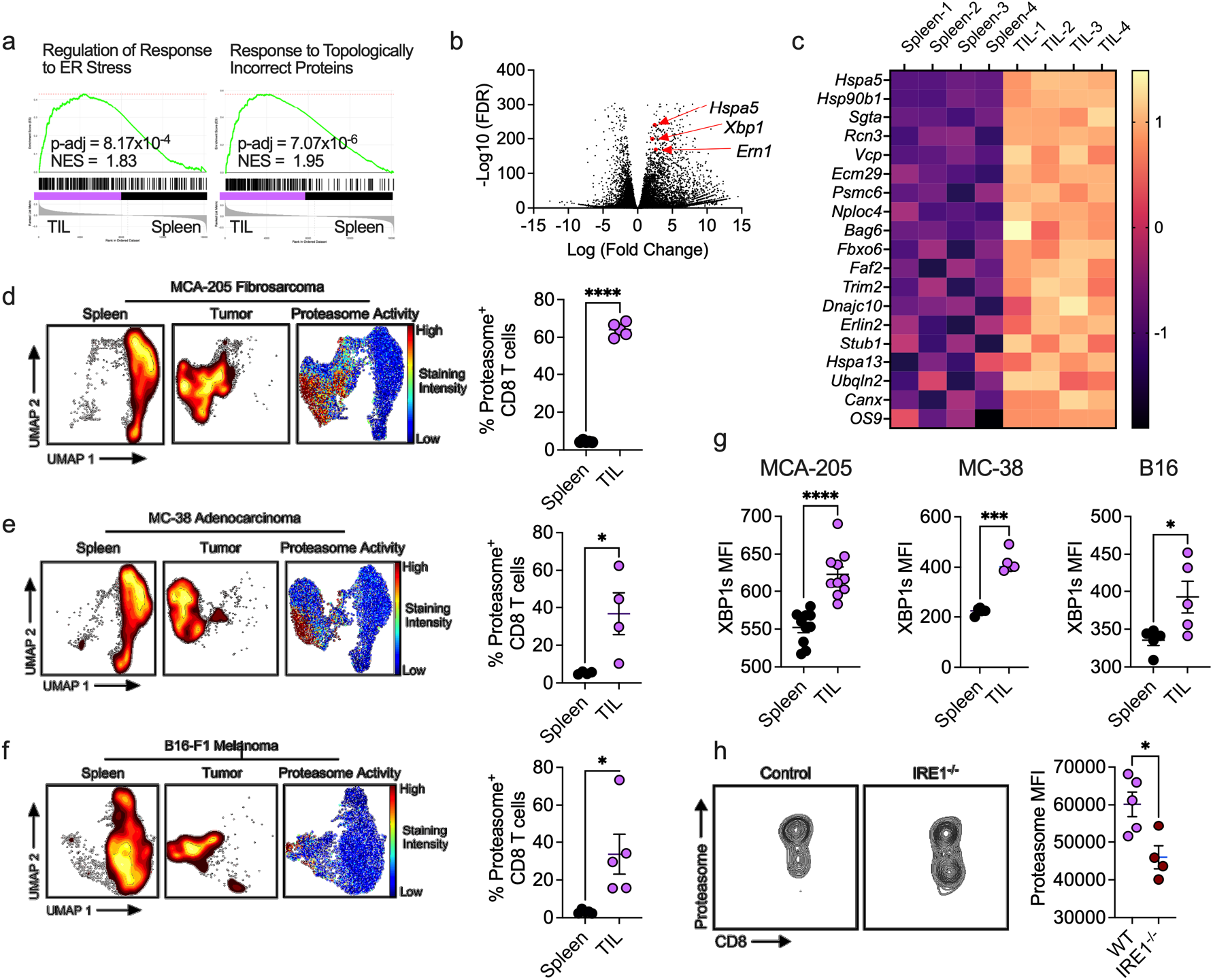
Proteasome activity identifies IRE1-regulated ER Stress in CD8^+^ TILs. a, Leading edge plots of GSEA and b, Volcano plot with IRE1-associated genes from RNA-seq data set of CD8^+^ T cells isolated from spleens or tumors of MCA-205 sarcoma bearing mice. c, ERAD pathway proteins identified by LFQ MS proteomics significantly regulated between CD8^+^ T cells isolated as in a. d-f, UMAP projection and quantification of Proteasome Probe MFI identified between spleen and TILs of indicated tumor models. g, XBP1s signaling in CD8^+^ T cell populations isolated as in d-f. h, Proteasome Probe MFI identified between WT and IRE1^−/−^ TILs 13-days post-infusion to B16 tumor-bearing hosts. Significance was calculated via a, Benjamini-Hochberg, d-h) unpaired student’s *t* test. * p < 0.05, ** p < 0.01, *** p < 0.001, **** p < 0.0001

A primary function of IRE1 signaling is to regulate the expression of proteins involved in ER-associated degradation (ERAD) – an ER proteostasis pathway involving the removal and proteasomal degradation of non-native proteins within the ER lumen^18^. Thus, we predicted that changes in IRE1-dependent regulation of ERAD could lead to alterations in the activity of specific aspects of this pathway, such as proteasome activity. We previously used a fluorescent probe to monitor proteasome activity in T cells^12^. Briefly, the probe contains a vinyl sulfone group that covalently interacts with the N-terminal threonine residue of all catalytic β-subunits of the proteasome enabling real-time identification of proteasome activity in live cells^19^. Through this work, we established increased proteasome activity as a biomarker of ER stress in CD8^+^ T cells^12^. Based on the known role of IRE1 in regulating ERAD, we reasoned that proteasome activity may allow for the identification of the CD8^+^ TIL compartment experiencing hyperactive IRE1 activity. In support of this notion, a proteomics screen identified proteins within the ERAD pathway as significantly enriched in CD8^+^ TILs isolated from MCA-205 sarcomas relative to autologous-matched CD8^+^ splenocytes (Figure 1c). We developed a high dimensional spectral flow cytometry panel specific for CD8^+^ T cells that included the proteasome activity probe in addition to markers of activation, differentiation, and dysfunction to monitor IRE1-regulated ER stress *in vivo*. Using the Uniform Manifold Approximation and Project (UMAP) dimension reduction algorithm, we visualized significant differences in the global phenotype of CD8^+^ T cells from the spleen and MCA-205 tumors. Projecting the fluorescent intensity of the proteasome probe onto the UMAP space, we found that the highest levels of proteasome activity were evident in CD8^+^ TILs (Figure 1d). We validated these data in MC-38 colon adenocarcinomas and B16-F1 melanomas (Figure 1e-f), establishing that enhanced protein degradation is a global trait of tumor-infiltrating CD8^+^ T cells.

Activation of IRE1 induces the unconventional splicing of mRNA encoding the transcription factor XBP1s, which can be monitored by intracellular flow cytometry. To determine whether proteasome activity correlated with IRE1 signaling, we assessed XBP1s expression among CD8^+^ T cells isolated from spleens and tumors in MCA-205, MC-38, and B16 tumors. In line with our RNA-sequencing and proteomics screens, we identified that XBP1s was uniformly increased in CD8^+^ TILs above autologous-matched splenic T cells across the three tumors models assessed (Figure 1g). To further evaluate the link between proteasome activity and IRE1 signaling in CD8^+^ TILs, we generated *IRE1^−/−^* T cells via CRISPR-Cas9 genome editing. OT-1 T cells were activated for 3 days followed by electroporation of guide RNAs specific for IRE1 or non-targeted controls. These cells were then allowed to differentiate for 4 days in IL-2 without stimulation. CD45.1^+^ mice bearing 7-day established B16-OVA tumors were then given IRE1^−/−^ or control T cells. 2-weeks later, the tumors were harvested, and the proteasome activity of adoptively transferred cells was assessed. We observed that IRE1^−/−^ T cells displayed a significant reduction in their proteasome activity when compared to the scramble control (Figure 1h). Together, our data show that proteasome activity in CD8^+^ TILs efficiently reports on IRE1 activity.

### ER Stress accumulates in dysfunctional CD8^+^ TILs

In the acute and adaptive phase of IRE1 signaling, the degradation of proteins through ERAD aims to reduce the irregular protein burden in the ER. However, hyperactive, chronic IRE1 activity impedes T cell tumor control^7, 8^. We reasoned that persistent IRE1 signaling could identify dysfunctional CD8^+^ TILs. Therefore, we assessed the temporal nature of proteasome activity to determine whether IRE1-regulated signaling in the context of T cells in tumors was persistent. We established cohorts of mice with MCA-205 fibrosarcoma every 72 hours, harvested tumors together after 14, 11, and 8 days of growth, then analyzed tumors using deep spectral flow cytometric phenotyping. Using the same high dimensional flow cytometry panel (Figure 1), we employed dimension reduction via the UMAP algorithm on tumor samples from all time points collected and then projected concatenated samples from their respective time points onto the UMAP space. First, we identified that CD8^+^ TILs acquired proteasome activity in a stepwise fashion over the course of tumor progression (Figure 2a). Additionally, we noted a significant alteration in CD8^+^ TIL phenotype across time illustrated along the UMAP dimension 1 that correlated with heightened proteasome activity in CD8^+^ TILs (Figure 2b). These data suggest that ER stress afflicts CD8^+^ TILs in a persistent manner that intensifies over time.

**Figure 2.**
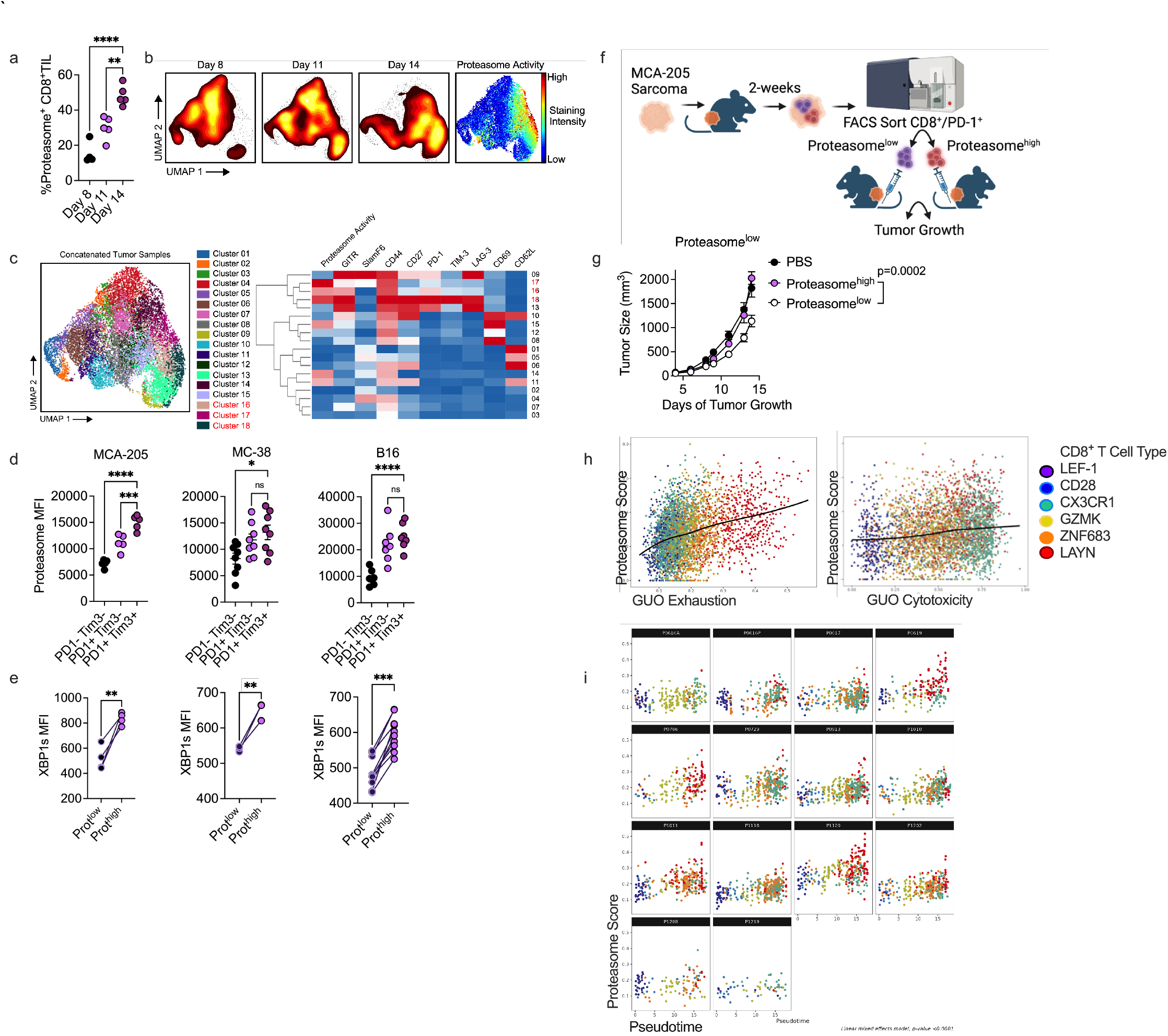
ER stress accumulates in dysfunctional CD8^+^ TILs. a, Quantification and b, UMAP projections of spectral flow cytometric data generated from proteasome monitoring in CD8^+^ TILs isolated from MCA-205 sarcomas across multiple time points post injection. c, Clustering analysis of deep phenotyping of samples collected in a-b. FACS phenotyping of d, proteasome activity and e, XBP1s expression in CD8^+^ TIL subsets isolated from indicated tumor models. f, Cartoon of experimental design and g, tumor control across time of PBS, Proteasome^high^, or Proteasome^low^ CD44^+^PD-1^+^/CD8^+^ MCA-205 sarcoma TILs infused into *Rag2^−/−^* mice bearing MCA-205 sarcomas. Data are representative of at least 2 independent experiments. Data points represent individual mice (n=5 mice per group). h, Compilation of correlation of KEGG Proteasome gene set expression and GUO Exhaustion score or GUO cytotoxicity score from n=3000 CD8^+^ T cells isolated from 14 lung cancer patient tumors, neighboring non-cancerous tissue, or PBMC with corresponding pseudotime clustering information^23^, i, individual patient signatures as in h. Significance was calculated via a,d, One way ANOVA with Dunnett’s multiple comparison test, e, paired student’s *t* test, g) simple linear regression, h-i) Spearman correlation. * p < 0.05, ** p < 0.01, *** p < 0.001, **** p < 0.0001

A hallmark of CD8^+^ T cell dysfunction in cancer is the progressive accumulation of inhibitory receptors and subsequent loss of function^20^. Given that our spectral flow cytometry panel includes several inhibitory receptors and markers of differentiation, we deepened our analysis by assessing the expression levels of these markers as they relate to proteasome activity in CD8^+^ TILs. We performed unsupervised clustering analysis on the viable CD8^+^ TIL pool using the FlowSOM algorithm from all the tumor samples across all time points to partition the data. This resulted in 18 unique clusters which we projected onto the UMAP space with mean expression levels of select markers hierarchically clustered into a heatmap (Figure 2c). Focusing on the highest relative levels of proteasome activity, clusters 16, 17, and 18 stood out (in red). In particular, cluster 18 displayed high levels of inhibitory receptors PD-1, Tim-3, and Lag-3, suggesting persistent proteasome activity was associated with T cell dysfunction. Using co-staining of PD-1 and Tim-3 as a surrogate for dysfunction^20^, we confirmed a stepwise progression in proteasome activity as T cells acquired hallmarks of dysfunction in multiple solid cancer types, indicating that proteasome activity is inexorably linked with progressive CD8^+^ T cell dysfunction (Figure 2d). These data were further complimented by our identification of increased XBP1s signaling in proteasome^high^ CD8^+^ TILs compared to proteasome^low^ counterparts in all three tumor models assessed (Figure 2e), linking hyperactive IRE1 signaling to overactivity of the proteasome.

Given that our phenotypic analysis of the proteasome active population correlated with the expression of markers of T cell dysfunction, we aimed to test the functional state of proteasome^high^ and proteasome^low^ CD8^+^ TIL populations. Reports have demonstrated the heterogeneous nature of CD8^+^ TIL-mediated tumor control based on phenotypic characteristics of the TIL pool^21, 22^. However, to this point, the cell-intrinsic role of ER stress to impede or augment TIL populations is poorly understood. To test the contribution of chronic proteasome activity to affect CD8^+^ TIL tumor control, we FACS sorted live CD45^+^ CD8^+^ CD44^+^ PD-1^+^ TILs from established MCA-205 fibrosarcomas based on proteasome expression levels and then adoptively transferred the TIL populations into *Rag2*^−/−^ mice bearing 2-day MCA-205 fibrosarcomas, and assessed tumor growth over time (Figure 2f). Mice receiving the proteasome^low^ CD8^+^ TILs showed significantly slower tumor progression compared to the proteasome^high^ CD8^+^ TILs and non-T cell treated controls (Figure 2g). Taken together, these data suggest proteasome^high^ CD8^+^ TILs exhibit poor antitumor function relative to proteasome^low^ counterparts.

The data to this point prompted us to study whether we could link increased proteasome activity to CD8^+^ TIL dysfunction in cancer patient samples. We identified a single cell RNA-seq data set from human lung cancer patients that effectively differentiated cytotoxic and exhausted T cells^23^. We first validated our efforts by reproducing the defined CD8^+^ T cell state clusters, C1-LEF-1 (Naïve), C2-CD28, C3-CX3CR1 (Cytotoxic), C4-GZMK, C5-ZNF683, and C6-LAYN (Exhausted), reported previously^23^. Moreover, we were able to recreate the pseudotime analysis that separated the naïve, cytotoxic, and exhausted populations (Supplemental Figure 1). Having validated the approach, we assessed proteasome activity in the comparative T cell states by generating a proteasome score using the KEGG Proteasome Pathway and projecting that against both the cytotoxicity and exhaustion pseudotime generated from Guo and colleagues^23^ (Figure 2h). We observed that as cells traveled down the exhaustion differentiation pathway, there was a significant upregulation of the proteasome score whereas cells in the cytotoxic branch demonstrated a more muted proteasome score (Figure 2h). When we partitioned the data across individual cancer patients, we found a similar trend where proteasome score was associated with the accumulation of exhausted CD8^+^ T cells (Figure 2i). These data demonstrate that hyperactive proteasome activity identifies dysfunctional CD8^+^ TILs in humans and mice.

### Ablation of IRE1 ameliorates hyperactive ER stress

T cell dysfunction in the solid TME is induced by multiple stressors including chronic antigen stimulation, nutrient deprivation^24, 25^, and hypoxia^26^. We previously observed that T cell receptor (TCR) stimulation engages the UPR^27^, and we have also found that exposure to the TME in the absence of TCR stimulation induces ER Stress^12^. To elucidate the biological requirements that drive persistent IRE1-associated ER stress *in vivo*, we studied proteasome activity in the context of chronic T cell response to virus and in response to the same viral antigen in the TME. We devised an experimental model where CD45.1/CD45.2^+^ P14 and CD45.2^+^ OT-1 transgenic T cells were co-transferred into CD45.1^+^ gp33-B16F1 melanoma-bearing mice followed by tumor harvest 2 weeks later. Simultaneously, in a separate cohort of mice we adoptively transferred CD45.1/CD45.2^+^ P14 T cells into naïve CD45.1^+^ mice 24 hours prior to challenge with LCMV-Clone 13 (LCMV-Cl13) followed by splenic harvest 14 days later (Figure 3a). This enabled us to assess whether chronic antigen stimulation (P14 from LCMV-Cl13 spleens), the TME (OT-1 from gp33-B16F1 tumors), or a combination of the two (P14 from gp33-B16F1 tumors) amplifies proteasome activity. First, antigen-specific P14 T cells harvested from the spleen of LCMV-Cl13 infected mice showed no differences in proteasome activity when compared to the endogenous T cell pool, suggesting chronic antigen stimulation alone did not induce IRE1-associated ER Stress (Figure 3b). While the magnitude of induction varied, TME exposure prompted robust proteasome activity in all T cell pools assessed including endogenous, non-tumor specific (OT-1), and tumor antigen-specific (P14) TILs compared to autologous splenic pools (Figures 3c-3d). Focusing specifically on CD8^+^ TILs, we found that P14 T cells harbored greater proteasome activity compared to OT-1 and endogenous TIL pools (Figure 3d). Taken together, these data suggest that antigen specificity in the context of the TME exacerbates ER stress in CD8^+^ TILs.

**Figure 3.**
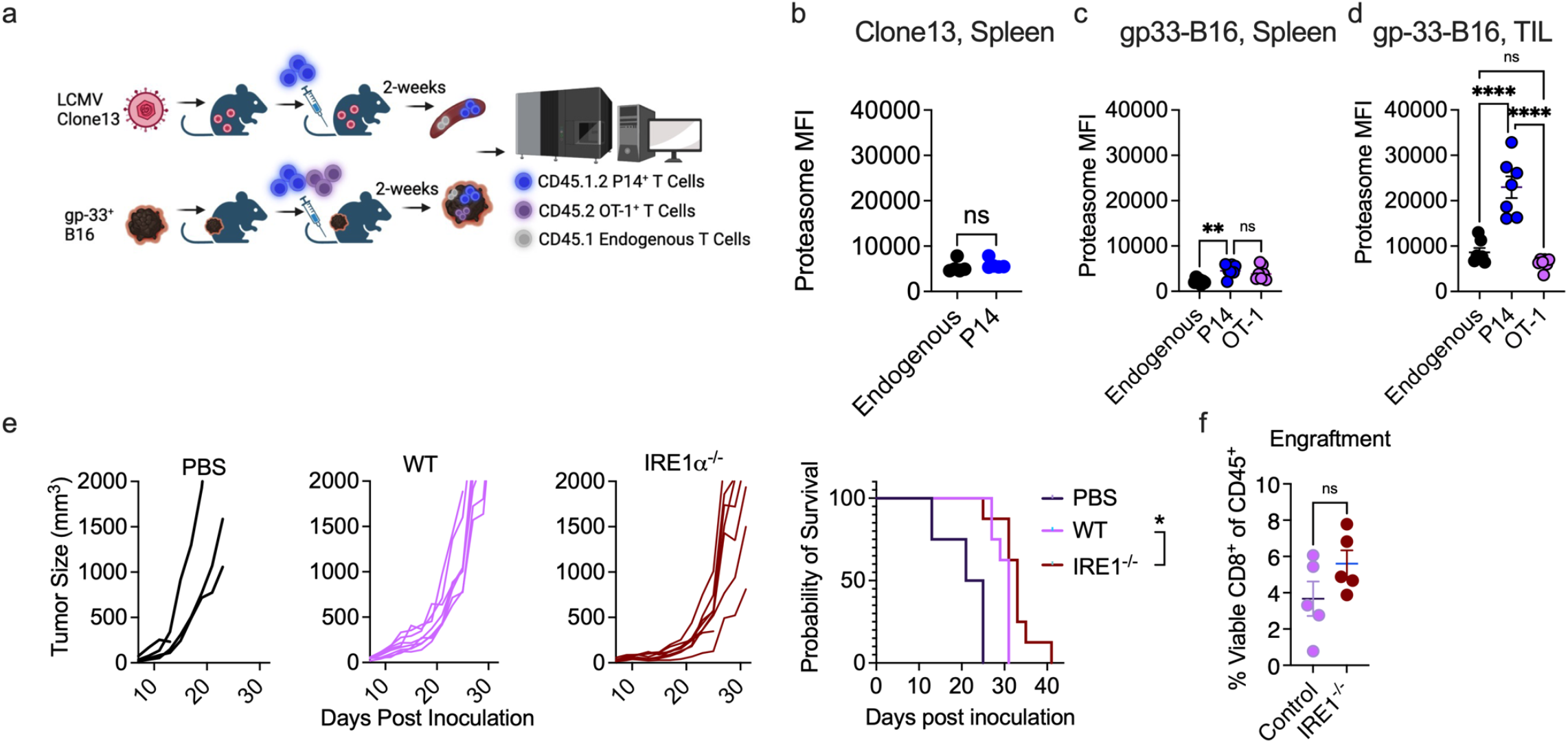
Ablation of IRE1 ameliorates hyperactive ER stress. a, Cartoon of experimental design and quantification of proteasome activity in endogenous, P14, and/or OT-1 T cells harvested from b, spleens of mice bearing chronic LCMV infection or c, spleens of mice bearing gp33-expressing B16 melanomas or d, tumors of mice bearing gp33-expressing B16 melanomas. Tumor growth and survival of mice bearing B16-OVA tumors infused with e, WT or IRE1^−/−^ OT-1 T cells or f, enumeration of engraftment 5-days post transfer of mice treated as in e. Significance was calculated via b-c, unpaired student’s *t* test, d, one-way ANOVA with a Tukey’s multiple comparison test, e-f, g, simple linear regression and long rank test. * p < 0.05, ** p < 0.01, *** p < 0.001, **** p < 0.0001

In cells the initial phase of the IRE1 response is protective, enabling cells under ER stress to adapt to adverse microenvironment conditions in response to perturbed ER homeostasis^28, 29^. In contrast, persistent IRE1 can result in hyperactive IRE1 signaling that promotes metabolic dysregulation and cell death^30–32^. Our data to this point illustrate that persistent antigen stimulation in the TME promoted hyperactive IRE1 signaling that restricts tumor control. To test that IRE1 signaling is deleterious to T cell-mediated tumor immunity, we generated IRE1^−/−^ T cells using the CRISPR-Cas9 approach outlined above. Indeed, cells treated with IRE1 guide RNAs were devoid of IRE1 expression (Supplemental Figure 2). While the adoptive transfer of IRE1 deficient tumor-specific OT-1 T cells resulted in increased tumor control and extended survival compared to IRE1 competent T cells, the effects only narrowly reached statistical significance (Figure 3e). To gain insight into the *in vivo* attributes of IRE1^−/−^ CD8^+^ TILs, we harvested tumors from animals infused with WT or IRE1^−/−^ CD8^+^ T cells and enumerated T cell engraftment as a measure of viable cells in tumors 5 days post transfer. As a potential explanation for limited benefits to tumor control, we observed that IRE1 deficiency did not confer enhanced engraftment of T cells relative to WT T cells (Figure 3f). Therefore, while loss of IRE1 limited hyperactive signaling evidenced by diminished proteasome activity (Figure 1), *in vivo* data suggested that stress sensor ablation was unable to confer protective effects to T cells during acute stage engraftment in the TME.

### The adaptive IRE1 UPR bolsters T cell metabolism and tumor control

IRE1 activators such as IXA4 have been developed to selectively enhance the adaptive and protective phase of the IRE1 UPR while limiting the deleterious effects associated with chronic IRE1 activation^33^. We reasoned that the ER environment of T cells could be enhanced through IXA4-dependent IRE1 activation, enabling heightened capacity to cope with misfolded/ damaged proteins in the TME. We activated T cells in the presence of IXA4 and confirmed that XBP1s was significantly increased in CD8^+^ T cells post-activation (Figure 4a). To determine whether IXA4 limited markers of hyperactive IRE1 signaling in T cells *in vitro*, we measured proteasome activity over the course of T cell expansion. We observed a peak in proteasome activity in T cells 4 hours after activation with IXA4; however, 24-, 48-, and 72-hours post-activation, proteasome activity was markedly reduced compared to vehicle controls (Figure 4b). These data suggest that IRE1 hyperactive signaling was limited by IXA4 treatment over the course of T cell expansion.

**Figure 4.**
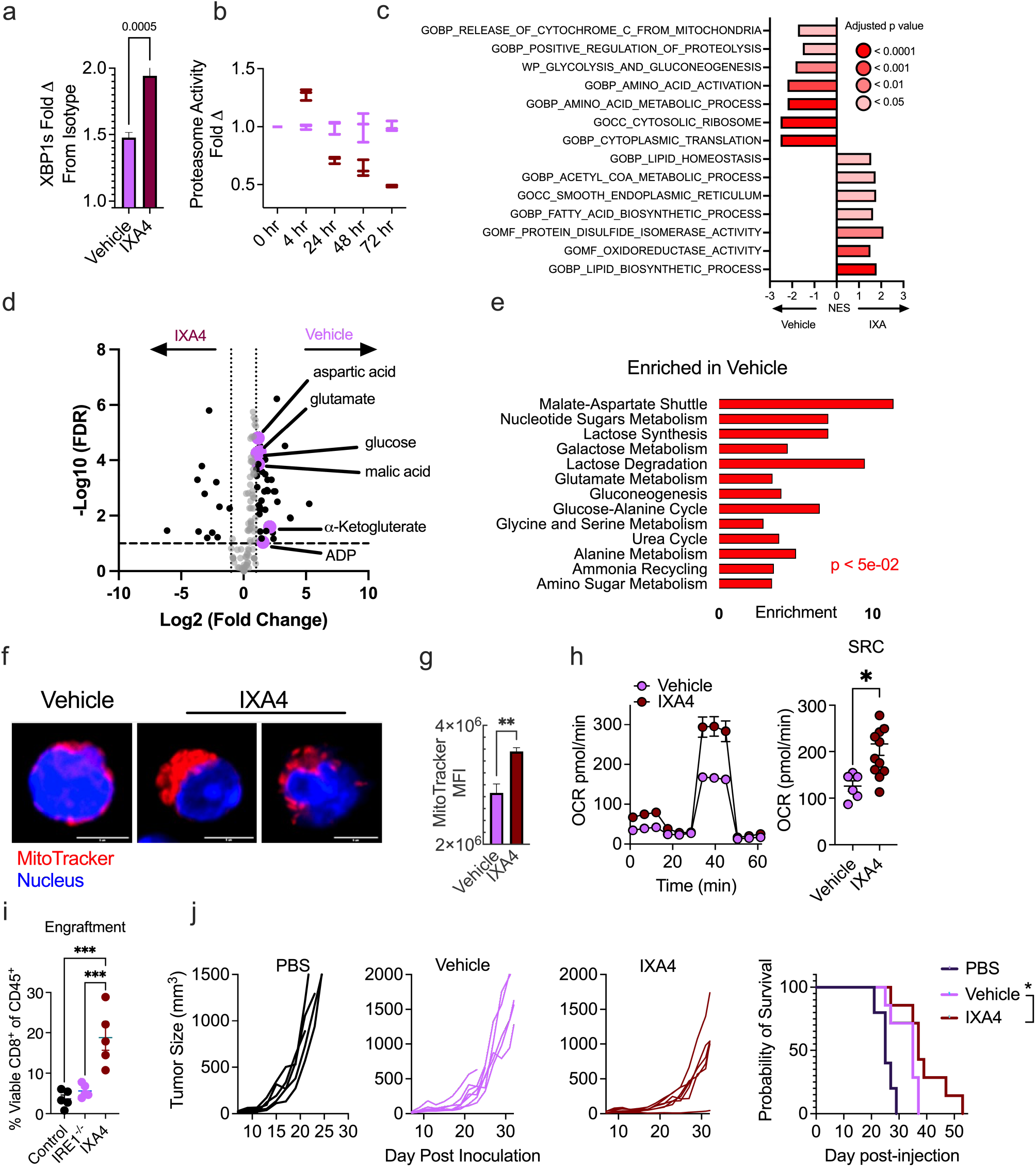
The adaptive IRE1 UPR bolsters T cell metabolism and tumor control. a, XBP1s expression and b, proteasome activity in vehicle and IXA4-treated T cells. c, Gene sets of interest enriched in indicated T cell groups identified by RNA-seq performed 4 hours after T cell activation. d, Volcano plot of significantly regulated metabolites enriched from LC/MS metabolomics with e, sequential quantitative enrichment analysis. f, confocal imaging, g, FACS quantification of MitoTracker and h, Seahorse Bioanalysis OCR trace and quantification of spare respiratory capacity performed on vehicle and IXA4-treated T cells. i, Engraftment 5 days post infusion of WT, IRE1^−/−^, or IXA4-treated T cells infused into mice bearing B16-OVA tumors, and j, tumor control and survival of OT-1 T cells treated with IXA4 then infused into animals bearing B16-OVA tumors. Significance was calculated using a-b, g-i) unpaired student’s *t* test, c) Benjamini-Hochberg correction, D) hypergeometric test, j) simple linear regression or Log Rank test. * p < 0.05, ** p < 0.01, *** p < 0.001, **** p < 0.0001.

Next, we performed RNA-sequencing to study gene programming in response to IXA4 treatment. We observed that IXA4 enforced transcriptional programming of ER protein folding enzymes, lipid metabolism, and ER structure and transport systems, consistent with the known role of IRE1/XBP1s in regulating these pathways^33^. Comparatively, vehicle control T cells upregulated pathways involved in protein translation, amino acid turnover, glycolytic metabolism, and mitochondrial dysfunction (Figure 4c). This suggested that IXA4 induced differential metabolic programming in T cells. To further distinguish between the differential metabolic signatures observed in vehicle and IXA4-conditioned T cells, we performed LC/MS metabolomics followed by sequential enrichment analysis. Strikingly, we observed metabolites and significantly upregulated metabolic pathways in vehicle-treated T cells that indicated the use of amino acids to fuel glucose production and energy synthesis. These data were in line with the known role of the ubiquitin-mediated proteasome system to catabolize proteins to amino acids that in turn serve as an energy resource^34^. Specifically, relative to IXA4-treated T cells, vehicle T cells upregulated glutamate, aspartic acid, α-ketoglutarate, malic acid, and glucose (Figure 4d). Enrichment analysis indicated that these metabolites were associated with increased activity of the malate-aspartate shuttle, which plays a key role in maintaining glycolysis through modulating the balance of energy carrier NAD+/NADH^35^. Moreover, α-ketoglutarate, glutamate, and glucose drove enrichment of the glucose-alanine cycle and gluconeogenesis which work in tandem to catabolize proteins to alanine that is then converted to free glucose^36^. Finally, vehicle T cells were enriched in multiple pathways associated with metabolism of amino acids and the byproducts of amino acid breakdown, including urea cycle and ammonia cycling^37^ (Figure 4e). Together, these data suggest that IXA4-treated T cells comparatively limit glucose production that is fueled by proteasome-driven recycling of amino acids.

Given these distinct metabolic profiles, as well as the downregulation of gene signatures associated with poor mitochondrial health in IXA4-treated T cells (Figure 4c), we next assessed mitochondrial health in vehicle or IXA4-treated T cell groups. Both confocal imaging and flow cytometry analysis using MitoTracker revealed that IXA4-primed T cells harbored increased mitochondrial mass (Figure 4f-g). Further, mitochondria in IXA4-treated T cells showed increased basal respiration, ATP-linked respiration, and spare respiratory capacity, as compared to vehicle-treated cells (Figure 4h), indicating that IXA4-treated cells harbored improved capacity to produce mitochondrial ATP^38^. In line with this, we observed that IXA4 treatment engendered a stem-like phenotype in T cells *in vitro* compared to vehicle controls (Supplemental Figure 3). This prompted us to infuse vehicle and IXA4-treated OT-1 T cells into tumor-bearing hosts to measure their capacity to control tumor growth. Consistent with the phenotypic and metabolic benefits conferred by IXA4 treatment, IXA4 treatment robustly increased T cell engraftment in host tumors relative to vehicle controls, an effect eclipsed in the engraftment observed in IRE1-deficient T cells (Figure 4i). In turn, we observed that mice infused with IXA4-treated T cells exhibited superior tumor control and enhanced survival relative to vehicle controls (Figure 4j). These data demonstrate the protective contributions of IRE1/XBP1s signaling to augment T cell tumor control.

## Discussion

As tumor-specific CD8^+^ T cells accumulate in the TME, they experience chronic, hyperactive IRE1 signaling. In response, TILs perpetually engage the proteasome to manage the perceived protein stress, an observation that enabled the identification of the CD8^+^ T cell subset afflicted by persistent IRE1 signaling. While abrogation of IRE1 was able to circumvent the harmful aspects of the chronic stress response, it also denied CD8^+^ TILs of the protective benefits of adaptive signaling. Transient activation of the IRE1 pathway via pharmacological stimulator IXA4 led to the remodeling of the CD8^+^ T cell transcriptome, metabolome, and phenotypic traits allowing for significantly enhanced tumor control. By preemptively inducing IRE1-signaling, we improved T cell capacity to manage the misfolded protein burden, averting toxic cellular states instilled by the TME. This indicates that pharmacologic enhancement of adaptive IRE1-XBP1s signaling in T cells prior to infusion delivers tangible benefits to CD8^+^ T cell-mediated immunotherapy.

The cellular response to ER stress can be partitioned into two distinct phases which are associated with divergent physiological outcomes^39, 40^. The adaptive UPR aims to enhance cell survival by initiating pathways that allow the cell to cope with and overcome ER stress^41^. Conversely, when cells are undergoing chronic, hyperactive signaling, the response shifts toward apoptotic cell death^40^. A significant barrier to monitoring phases of the UPR in complex systems, such as the tumor microenvironment, is the lack of conclusive techniques to assess the extent of ER stress. By linking proteasome activity to IRE1 activity, we were able to overcome this obstacle to provide a key marker of T cell exhaustion and demonstrate that the TME induces chronic ER stress. Using sequential analysis of TIL populations in early and late-stage tumors, we show that ER stress is unresolved in CD8^+^ TILs, evidenced by high proteasome activity and chronic IRE1 activation. Exacerbation of ER stress as T cells differentiate and age in tumors suggests that CD8^+^ TILs reach a critical juncture, specific to the TME, at which the protein burden can no longer be managed, committing CD8^+^ TILs to dysfunction and thereby serving as an intrinsic checkpoint to T cell-mediated tumor clearance. To this end, recent work has identified a previously undiscovered T cell subset specific to cancer immunotherapy resistance that is defined by its stress response state, further supporting our notion of the UPR mediated checkpoint^4^. As current interventional strategies targeting IRE1 have yet to deliver meaningful results^42^, our data offer clues as to why. While mitigating the chronic UPR signaling provides some benefit^13^, total ablation dually eliminates the protective effects of adaptive signaling. As such, identifying patients most likely to respond to immunotherapies may be intimately tied to the extent of hyperactive cell stress in individual patient TILs. To that end, the extreme heterogeneity in the proteasome signature in both humans and mice suggests that T cells experience diverse levels of ER stress. It is interesting to speculate that patients with a high irregular protein burden may benefit from complete stress sensor targeting; whereas patients with limited damage to the cellular proteome may benefit from protective and transient UPR activation.

Our findings using selective IRE1-XBP1s activator IXA4 posit that the adaptive IRE1 UPR regulates T cell metabolic programming. Consistent with existing research on IRE1-XBP1s signaling, our study demonstrated that IXA4 treatment not only transcriptionally modulated ER function, but also altered the expression of genes involved in T cell lipid metabolism and homeostasis^43^. This IXA4-driven lipid remodeling was first observed in hepatic cells where XBP1s induction limited expression of certain lipogenic genes and decreased triglyceride content to mitigate liver steatosis^16^. Given that our metabolomics approach targeted polar metabolites, a nonpolar extraction method could enable further characterization of small non-polar lipid-associated metabolites likely enriched in IXA4-treated T cells. In hepatic cells, IXA4 further influenced metabolism by restraining gluconeogenesis and improving glucose homeostasis^33^. Similarly, our results indicated that IXA4 treatment redirected T cell metabolism away from glycolysis by reducing excessive glucose production, specifically from the catabolism of glucogenic amino acids. Since glycolysis is a metabolic niche already occupied and competed for by highly proliferative tumor cells, IXA4 metabolic reprogramming could better equip TILs to overcome the limited availability of glucose in the TME ^25^. Another metabolic obstacle TILs face in the TME is mitochondrial dysfunction characterized by small, fragmented, and hyperpolarized mitochondria^44^. We observed that IXA4 conditioning endowed T cells with increased mitochondrial mass and elevated mitochondrial ATP reserves. Thus, adaptive IRE1 signaling exhibits an ability to anticipate and circumvent the metabolic challenges unique to the solid tumor. As the cancer immunotherapy field endeavors to improve current cellular therapies in solid cancers, our work emphasizes the previously unidentified role of the adaptive and protective phase of the UPR in bolstering T cell metabolic fitness to cope with the bioenergetic constraints of the TME.

## Methods

### Lead contact

Further information and requests for resources and reagents should be directed to the lead contact Jessica Thaxton.

### Materials availability

This study did not generate new unique reagents.

### Data and code availability

All data reported in this paper will be shared by the lead contact upon request.

### Mice

OT-1 (C57BL/6-Tg(TcraTcrb)1100Mjb/J), P14 (B6.Cg-Tcra*tm1Mom*Tg(TcrLCMV)327Sdz/TacMmjax), CD45.1 (B6.SJL-*PtprcaPepcb*/BoyJ), and C57BL/6 mice were obtained from the Jackson Laboratories. All animal experiments were performed in accordance with protocol 21-228 approved by the Institutional Animal Care and Use Committee (IACUC) at the University of North Carolina at Chapel Hill-CH. Mice were housed in a pathogen-free animal facility and maintained by the Division of Comparative Medicine at UNC-CH. Age-matched (6-8 weeks) female mice were used in all mouse experiments. The number of animals (biological replicates) is indicated in figure legends.

### Cell Cultures

MCA-205 (Sigma), B16F1 (ATCC), B16F1-OVA (kind gift of Dr. Mark Rubinstein), and gp33-B16F1 (Kind gift of Dr. Justin Milner) tumor lines, as well as OT-1 and P14 T cells, were maintained in RPMI supplemented with 10% FBS, 300 mg/L L-glutamine, 100 units/mL penicillin, 100 µg/mL streptomycin, 1mM sodium pyruvate, 100µM NEAA, 1mM HEPES, 55µM 2-mercaptoethanol, and 0.2% Plasmocin mycoplasma prophylactic (InvivoGen). 0.8 mg/mL Geneticin selective antibiotic (Invitrogen) was added to media of B16F1-OVA and gp33-B16F1 cells for multiple passages then cells were passaged once in Geneticin-free media prior to tumor implantation. MC-38 (Kerafast) were cultured in DMEM supplemented with 10% FBS, 2mM glutamine, 0.1mM nonessential amino acids, 1mM sodium pyruvate, 10mM Hepes, 50ug/mL gentamycin sulfate, 100 units/mL penicillin, 100 µg/mL streptomycin, and 0.2% Plasmocin mycoplasma prophylactic (InvivoGen). MCA-205, MC-38, B16F1, B16F1-OVA, and gp33-B16F1 cell lines were determined to be mycoplasma-free in March of 2023. For OT-1 T cell activation and expansion, whole splenocytes from OT-1 mice were activated with 1 µg/mL OVA 257-264 peptide (InvivoGen) and expanded for 3 days with 200 U/mL rhIL-2 (NCI) then split and expanded for 4 more days. In some cases, IXA4 (30µM) was added at T cell activation and again at the cell split for 4 more days. For P14 T cell activation, CD8^+^ T cells were purified from P14 spleens using EasySep Mouse CD8^+^ T Cell Isolation Kit (Stemcell Technologies). Cells were then expanded using plate-bound CD3/CD28 (Biolegend) antibodies in the presence of 200 U/mL rhIL-2 (NCI) for 3-4 days prior to adoptive transfer.

### CRISPR-Cas9

IRE1 deficient OT-1 T cells (IRE1^−/−^) and non-targeting control T cells (sgNT control) were generated using the CRISPR-Cas9 system. For IRE1^−/−^ experiments, whole splenocytes from OT-1 mice were electroporated on day 3 after initial activation with 1 µg/mL OVA 257-264 peptide (InvivoGen), and cells were further expanded for 4 days with 200 U/mL rhIL-2 (NCI). Electroporation was performed with sgRNA/Cas9 RNP complexes using Alt-R Sp Cas9 Nuclease V3 (IDT) and using the 4D-Nucleofector with P3 Primary Cell 4D Nucleofector X Kit S (Lonza Bioscience).

### Seahorse Bioanalysis

Seahorse XF Mito Stress Tests and Seahorse Real-Time ATP Rate Assays were performed using the Seahorse XFe96 Analyzer. 96-well Seahorse plates (Agilent) were coated with CellTak (Corning). T cells were plated in Seahorse XF DMEM Medium (Agilent) supplemented with 1% fetal bovine serum (FBS) and centrifuged for adherence. For the Seahorse Mito Stress Test, 1µM oligomycin, 1.5mM FCCP, and 2µM rotenone/1µM antimycin A (Sigma) were injected sequentially, and the oxygen consumption rate (OCR) was measured. FCCP is used to elicit maximal respiration, however FCCP can inhibit OCR at high concentrations. An FCCP dose of 1.5 mM was selected based on prior optimization during performance of the Seahorse XF Cell Mito Stress test with 5 different doses of FCCP ^6, 45^. For Seahorse normalizations, equal numbers of T cells per condition were seeded on Cell Tak-coated plates (Agilent) immediately prior to performing the assay and cell counts were verified via microscopy directly after assay performance. Spare respiratory capacity was calculated as the difference between the basal and maximal OCR readings after addition of FCCP.

### Metabolomics

*Sample Preparation:* T cells were harvested and washed with sodium chloride solution then resuspended in 80% methanol for three freeze-thaw cycles at −80°C and stored at −80°C overnight to precipitate hydrophilic metabolites. Samples were centrifuged at 20,000xg and methanol was extracted for metabolite analysis. Remaining protein pellets were dissolved with 8M urea and total protein was quantified using Pierce BCA assay (Thermo Fisher). Total protein amount was used for equivalent loading for High Performance Liquid Chromatography and High-Resolution Mass Spectrometry and Tandem Mass Spectrometry (HPLC-MS/MS) analysis. *Data Acquisition*: Samples were dried with a SpeedVac then 50% acetonitrile was added for reconstitution followed by overtaxing for 30 sec. Sample solutions were centrifuged and supernatant was collected and analyzed by High-Performance Liquid Chromatography and High-Resolution Mass Spectrometry and Tandem Mass Spectrometry (HPLC-MS/MS). The system consists of a Thermo Q-Exactive in line with an electrospray source and an Ultimate3000 (Thermo Fisher) series HPLC consisting of a binary pump, degasser, and auto-sampler outfitted with a Xbridge Amide column (Waters; dimensions of 2.3 mm × 100 mm and a 3.5 µm particle size). The mobile phase A contained 95% (vol/vol) water, 5% (vol/vol) acetonitrile, 10 mM ammonium hydroxide, 10 mM ammonium acetate, pH = 9.0; B with 100% Acetonitrile. For gradient: 0 min, 15% A; 2.5 min, 30% A; 7 min, 43% A; 16 min, 62% A; 16.1-18 min, 75% A; 18-25 min, 15% A with flow rate of 150 μL/min. A capillary of the ESI source was set to 275 °C, with sheath gas at 35 arbitrary units, auxiliary gas at 5 arbitrary units and the spray voltage at 4.0 kV. In positive/negative polarity switching mode, an *m*/*z* scan range from 60 to 900 was chosen and MS1 data was collected at a resolution of 70,000. The automatic gain control (AGC) target was set at 1 × 10^6^ and the maximum injection time was 200 ms. The top 5 precursor ions were subsequently fragmented, in a data-dependent manner, using the higher energy collisional dissociation (HCD) cell set to 30% normalized collision energy in MS2 at a resolution power of 17,500. Besides matching m/z, metabolites were identified by matching either retention time with analytical standards and/or MS2 fragmentation pattern. All standards were purchased from either Sigma, Fisher Scientific or VWR. Metabolite acquisition and identification was carried out by Xcalibur 4.1 software and Tracefinder 4.1 software, respectively. *Statistical Analysis:* Post identification, samples were normalized by taking the peak area under the curve for each metabolite per sample and dividing by the quotient of the total ion count per sample over the lowest total ion count in the batch. Subsequent transformation of normalized data was carried out with auto scaling to account for heteroscedasticity (van den Berg et al., 2006). Metabolites that were below detection in all samples were removed from analysis; missing values were imputed with 1/5 of the minimum positive value of their corresponding variable. Differential metabolite abundance between groups of interest was defined as absolute fold change > 1.5 and FDRq Adj p-value < 0.05. Quantitative enrichment analysis was performed using centered and scaled levels of all detectable metabolites after removal of invariant metabolites using the SMPDB database to identify biological processes associated with differential expression. All statistical analysis was performed using the MetaboAnalyst 5.0 web server (Pang et al., 2021).

### Proteomics

*Sample Preparation:* T cells were washed in PBS and lysed in 9M urea, 50 mM Tris pH 8, and 100 units/mL Pierce Universal Nuclease (Thermo Fisher Scientific) and the concentration of protein was measured using a Pierce BCA Assay (Thermo Fisher Scientific). Protein was trypsin (Sigma Aldrich) digested at 37°C for 18 hours, and the resulting peptides were desalted using C18 ZipTips (Millipore). *LC/MS data acquisition parameters:* Peptides were separated and analyzed on an EASY nLC 1200 System (Thermo Fisher Scientific) in-line with the Orbitrap Fusion Lumos Tribrid Mass Spectrometer (Thermo Fisher Scientific) with instrument control software v. 4.2.28.14. Peptides were pressure loaded at 1,180 bar, and separated on a C18 Reversed Phase Column [Acclaim PepMap RSLC, 75 μm x 50 cm (C18, 2 μm, 100 Å); Thermo Fisher Scientific] using a gradient of 2%–35% B in 120 minutes (Solvent A: 0.1% FA; Solvent B: 80% ACN/0.1% FA) at a flow rate of 300 nL/minute at 45°C. Mass spectra were acquired in data-dependent mode with a high resolution (60,000) FTMS survey scan, mass range of m/z 375–1,575, followed by tandem mass spectra of the most intense precursors with a cycle time of 3 seconds. The automatic gain control target value was 4.0e5 for the survey MS scan. Fragmentation was performed with a precursor isolation window of 1.6 m/z, a maximum injection time of 50 ms, and HCD collision energy of 35%; the fragments were detected in the Orbitrap at a 15,000 resolution. Monoisotopic precursor selection was set to “peptide.” Apex detection was not enabled. Precursors were dynamically excluded from resequencing for 20 seconds and a mass tolerance of 10 ppm. Advanced peak determination was not enabled. Precursor ions with charge states that were undetermined, 1, or >7 were excluded. *Mass spectrometry data processing:* Protein identification and quantification were extracted from raw LC/MS-MS data using the MaxQuant platform v.1.6.3.3 with the Andromeda database searching algorithm and label free quantification (LFQ) algorithm. Data were searched against a mouse Uniprot reference database UP000000589 with 54,425 proteins (March, 2019) and a database of common contaminants. The FDR, determined using a reversed database strategy, was set at <1% at the protein and peptide level. Fully tryptic peptides with a minimum of seven residues were required including cleavage between lysine and proline. Two missed cleavages were permitted. LC/MS-MS analyses were performed in triplicate for each biological replicate with match between runs enabled. The “fast LFQ” was disabled and “stabilize large ratios” features were enabled. The first search was performed with a 20 ppm mass tolerance, after recalibration a 4.5 ppm tolerance was used for the main search. A minimum ratio count of 2 was required for protein quantification with at least one unique peptide. Parameters included static modification of cysteine with carbamidomethyl and variable N-terminal acetylation. The protein groups’ text file from MaxQuant was processed in Perseus v. 1.6.5.0. Identified proteins were filtered to remove proteins only identified by a modified peptide, matches to the reversed database, and potential contaminants. Normalized LFQ intensities were log2 transformed. The LFQ intensities of technical replicates were averaged for each biological replicate. Quantitative measurements were required in at least three of five biological replicates in each treatment group. *Statistical Analysis:* Log2 transformed, protein LFQ intensities, normalized in MaxQuant, exhibited normal distributions. Binary comparisons of each LC/MS-MS analysis yielded Pearson correlation coefficients > 0.90 between all technical and biological replicates within each experiment. For comparison of spleen and tumor T cells, quantitative measurements were required in at least three of five biological replicates in each group. With a permutation-based FDR of 5% and S0 = 0.01, q val < 0.05, 177 proteins were significantly different between spleen and tumor T cells. toppgene.cchmc.org Toppfun software package was used to measure significantly enriched GO:BP pathways.

### Microscopy

For MitoTracker labeling, cells were incubated in 500nM MitoTracker Deep Red FM (ThermoFisher) for 30 minutes at 37°C and washed twice in PBS. Cells were fixed at room temperature in 4% paraformaldehyde in PBS for 15 minutes. Cells were washed 1x with PBS and incubated with nuclear stain DRAQ5 (Abcam) in PBS (1:1000) for 15 minutes, followed by 2x washes with PBS. A 96-well optical glass bottom plate image plate (Cellvis) was coated with CellTak (Corning), washed with DI water, and air dried. Labeled cells were adhered to the imaging plate surface by centrifugation for 2 minutes at 450 x *g*. Cells were imaged using the Zeiss LSM-880 confocal microscope with ZEN acquisition software (Zeiss) and processed with Bioimage ICY (v2.4.3.0)^46^ with the “Scale Bar” (Thomas Provoost, Pasteur Institute; http://icy.bioimageanalysis.org/plugin/scale-bar/) (v3.3.1.0) plug-in.

### Flow Cytometry

For high dimensional T cell phenotyping, cells were stained for 30 minutes at 4°C with a mix of viability dye and extracellular antibodies followed by overnight fixation using the Foxp3/Transcription Factor Fixation/Permeabilization Concentrate Diluent kit and Permeabilization Buffer (eBioscience). Cells were then washed with 1x permeabilization buffer before intracellular staining at room temperature for 3-4 hours. Samples were collected on a Cytek Northern Lights or in some cases on a BD Accuri C6 Plus. For all spectral flow cytometry experiments, single stain compensations were created using UltraComp eBeads Plus Compensation Beads (Invitrogen) or heat-killed cell samples. Extracellular antibodies used were some variation of the following: Zombie NIR viability dye (Biolegend), CD45-BV510 (BD Biosciences), CD45.2-BV510 (Biolegend), CD45.1-PE-Cy7 (Invitrogen), CD4-APC Fire 810 (Biolegend), CD11b-Alexa Fluor 532 (Invitrogen), CD8-Spark NIR 685 (Biolegend), TCRβ BV570, CD44-Brilliant Violet 785 (Biolegend), CD62L-BV421 (BD Biosciences), PD-1-BB700 (BD Biosciences), TIM-3-BV711 (Biolegend), LAG-3-APC-eFluor 780 (Invitrogen), SlamF6-APC (Invitrogen), CD127-Superbright 436 (eBioscience), KLRG1-Pacific Orange (Invitrogen), CD69-PE-Cy5 (Biolegend), GITR-BV650 (BD Biosciences), CD27-BV750 (BD Biosciences). Intracellular antibodies: CTLA-4-PE-Dazzle 594 (Biolegend), Ki67-PerCP-eFluor 710 (Invitrogen), TOX-PE (Miltenyi Biotec), TCF1/7-PE-Cy7 (Cell Signaling Technology), Granzyme B-Alexa Fluor 700 (Biolegend), and BCL-2-Alexa Fluor 647 (Biolegend). For experiments using the Proteasome Activity Probe (R&D), samples were counted and plated at equal numbers before being stained for 1 hour 45 minutes at 37°C with 1.25μM of the probe in PBS. Cells were then washed and stained with extracellular antibodies as described above before being fixed for 1 hour at room temperature. Samples were then acquired using the Cytek Northern Lights as described above. For XBP1s experiments, samples were plated and stained with some variation of CD8-FITC (Invitrogen), Live or Dye-PE (Biotium), and CD45.2-PE-Cy7 (Invitrogen) prior to being fixed overnight using the FoxP3 transcription factor staining kit. Cells were then washed with 1x permeabilization buffer prior to staining with XBP1s-AlexaFluor 647 for 3 hours at room temperature. Samples were acquired using the Accuri Platform (BD). For MitoTracker experiments, cells were incubated in 500nM MitoTracker Deep Red FM (ThermoFisher) for 30 minutes at 37°C and washed twice in PBS. Cells were then stained with CD8-FITC and Live or Dye-PE in PBS for 30 minutes at 4°C and then collected on the Accuri.

High-dimensional flow cytometry analysis was performed using the Omiq.ai platform. Briefly, live CD8^+^ CD11b^-^ singlets were gated and subsampled to 1500 events per sample, or in some cases the maximum number of events possible for the gate. To facilitate the visualization of marker expression patterns, cells were mapped to a two-dimensional embedding using the Uniform Manifold Approximation and Projection (UMAP) algorithm using the standard settings generated by the OMIQ platform. Mean fluorescent intensities of select markers were then projected onto the UMAP space to further visualize the distinct phenotypes. For clustering analysis, the FlowSOM algorithm was used by selecting prespecified expression markers that had not been used in the gating strategy to arrive at the population of interest. The number of clusters was generated using an elbow-guided *k* approach. The corresponding clusters were then projected back on the UMAP space for visualization of distribution. Finally, hierarchical clustered heatmap was generated by concatenating all samples to examine the median marker intensity across all clusters generated from FlowSOM. In some cases, mean fluorescent intensity was generated using the FlowJo software platform.

### RNA-sequencing

Three OT-1 spleens were collected, and cells were activated in the same manner as described above in the presence of 30µM IXA4 or vehicle control for 4 hours. Cells were washed in PBS and pellets were flash frozen at −80°C. Samples were sent to the UNC Lineberger Comprehensive Cancer Center Translational Genomics Lab (TGL) for RNA isolation using a Thermo Scientific KingFisher Flex (Cat 5400610) and the Applied Biosystems MagMAX mirVana Total RNA Isolation Kit (A27828) according to the manufacturer’s protocol (MAN0011138). Up to 1 µg of total RNA was used for the construction of libraries using a Hamilton NGS STAR automated liquid hander and the Illumina TruSeq Stranded mRNA (Cat 20020594) kit according to the manufacturer’s protocol (1000000040498). Dual-indexed libraries were pooled and sequenced on an Illumina NextSeq2000 system (Cat 20038897) P2 flow cell (Cat 20046811) to a depth of ∼65 million clusters per library with a 2×50 bp paired-end configuration. *Data Processing & Analysis:* Raw fastq files were aligned to the GRCm38v100 version of the mouse genome and transcriptome using STAR 2.7.6a. Quantification of gene abundance for each sample was done by Salmon v1.4.0 and genes were kept for subsequent analysis if it contained more than 10 reads in at least one sample. Normalization and differential gene expression were performed using the DESeq2 1.22.2 Bioconductor package in R. The false discovery rate was used to correct for multiple hypothesis testing. Gene set enrichment analysis was performed using the FGSEA Bioconductor package to identify gene ontology terms and pathways associated with altered gene expression for each of the comparisons performed. Gene sets were obtained from the ‘Hallmark Pathways’, ‘Curated Pathways’ and ‘Gene Ontologies’ sets from MSigDB.

### Tumor Modeling

MCA-205 fibrosarcomas, MC-38 colon adenocarcinomas, and B16F1 melanomas were established subcutaneously by injecting 2.5 × 10^5^, 1 × 10^6^, and 2.5 x10^5^ cells into the right flank of C57BL/6 mice, respectively. After 8-14 days of tumor growth, spleens and tumors from groups of mice were harvested and tumors were processed using the Mouse Tumor Dissociation Kit and gentleMACS dissociator (Miltenyi Biotec) according to manufacturer’s protocol. For some experiments, samples were pre-enriched using EasySep Mouse CD8^+^ T Cell Isolation Kit (Stemcell Technologies) according to manufacturer’s protocol and sorted using the FACSAria II cell sorter. CD8^+^ spleen and pooled TIL samples were washed in PBS and frozen for RNA-seq and proteomic analysis.

To measure tumor growth mediated by proteasome TIL subsets, MCA-205 fibrosarcomas were established as described above. After 12 days of tumor growth, tumors from groups of mice were harvested, processed, pre-enriched, then PD-1/CD8^+^ Proteasome^high^ and Proteaseome^low^ TIL subsets were sorted using the FACSAria II cell sorter then 2.5 × 10^4^ TILs were infused in 100 µL PBS via tail vein into mice bearing 2-day established MCA-205 fibrosarcomas, and tumor growth was measured every other day with calipers.

### Single-cell processing & clustering

Raw single-cell count data and cell annotation data was downloaded from NCBI GEO (GSE99254)1. Count data was normalized and transformed by derivation of the residuals from a regularized negative binomial regression model for each gene2 (SCT normalization method in Seurat3, v4.1.1), with 5000 variable features retained for downstream dimension reduction techniques. Integration of data was performed on the patient level with Canonical Correlation Analysis as the dimension reduction technique4. Principal component analysis and UMAP dimension reduction were performed, with the first 50 PCs utilized in UMAP generation. Cells were clustered utilizing the Louvain algorithm with multi-level refinement. The data was subset to CD8^+^ T cells, which were identified utilizing the labels provided by Guo et al1. Cell type labels were confirmed via 1) SingleR5 (v1.8.1) annotation utilizing the ImmGen6 database obtained via celda(1.10), 2) cluster marker identification, and 3) cell type annotation with the ProjecTILs T cell atlas7 (v2.2.1). After subsetting to CD8^+^ T cells, cells were again normalized utilizing SCT normalization, with 3000 variable features retained for dimension reduction. Due to the low number of cells on per-patient level, Harmony8(v1.0) was utilized to integrate the data at the patient level, rather than Seurat. PCA and UMAP dimension reduction were performed as above.

### Proteasome score analysis

Proteasome scores were calculated via UCell, utilizing the genes in the KEGG_PROTEASOME gene set. Moran’s I test with the pseudotime trajectory graph as input was utilized to test spatial correlation of proteasome genes with pseudotime, utilizing all samples and cells. To assess differences in proteasome scores across pseudotime and CD8^+^ T cell types, linear mixed effects models were implemented, with CD8^+^ cell type (or pseudotime), tissue source (tumor, peripheral blood, and adjacent normal) as fixed effects and patient ID as a random effect with random uncorrelated slope and intercept. Only patients with all tissue types were included. The p-values for fixed effects were approximated via Satterthwaite’s method in the afex package.

### Adoptive Cellular Transfer

For adoptive cellular therapy experiments, B16F1-OVA or B16-gp33 melanomas were established subcutaneously by injecting 2.5 × 10^5^ cells into the right flank of C57BL/6 mice, and tumor-bearing hosts were irradiated with 5 Gy 24 hours prior to T cell transfer. After 7 days of tumor growth, 3-5 × 10^5^ OT-1 T cells conditioned with vehicle or 30µM IXA4 were infused in 100 µL PBS via tail vein into tumor-bearing mice. In some cases, OT-1 T cells treated with sg control or IRE1 RNPs were infused into tumor-bearing mice. Tumor growth was measured every other day with calipers, and survival was monitored with an experimental endpoint of tumor growth ≥ 300 mm^2^. For tumor harvest experiments, 1-2 x10^6^ of a given T cell group with respective controls were infused into B16-OVA tumor bearing mice 7-10 days after tumor challenge, and tumors were harvested 5-13 days later using the Mouse Tumor Dissociation Kit, and gentleMACS Octo dissociator with heaters (Miltenyi Biotec) according to the manufacturer’s protocol.

For LCMV experiments, naïve CD45.1/CD45.2^+^ P14^+^ CD8^+^ T cells were purified from spleens using the StemCell CD8^+^ isolation kit (STEMCELL Technologies) and 500 were adoptively transferred into CD45.1^+^ mice. 24 hours later, mice were challenged with 2.0 × 10^6^ PFU of LCMV-Clone 13. After 14 days of incubation, spleens of infected animals were harvested, and proteasome activity was assessed as described above.

### Statistical Analysis

GraphPad Prism v 9.3.0 was used to calculate p values using statistical tests as indicated in figure legends. Values of p < 0.05 were considered significant.

## Supplemental Information

Supplemental Information can be found in the Supplemental Information document.

## Author Contributions

Conceptualization, B.P.R., E.J.G., J.E.T., Methodology, B.P.R., E.J.G., J.J.M., P.G., J.E.T., Software, J.M., Validation, B.P.R. and E.J.G., Formal Analysis, B.P.R., E.J.G., A.S.K., E.G.H., K.E.H., E.L.S., Investigation, B.P.R., E.J.G., A.S.K., E.G.H., K.E.H., E.L.S., J.M, Resources, P.G., J.M.G., J.J.M., R.L.W., J.E.T., Writing – Original Draft, B.P.R., E.J.G., J.E.T., Writing – Review and Editing B.P.R., E.J.G., M.J.E., R.L.W., J.J.M., J.E.T. Visualization, B.P.R., E.J.G., J.M., J.E.T., Supervision, J.E.T., Funding Acquisition, J.E.T.

