## Supplemental Figures for "Adaptive IRE1 Signaling Elicits T Cell Metabolic Remodeling and Tumor Control"

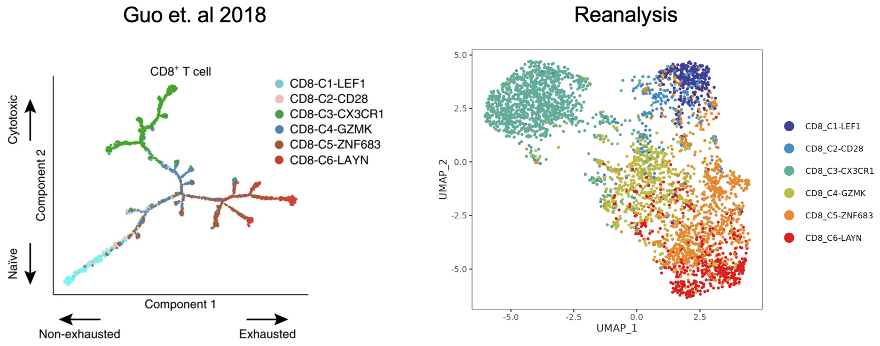


**Supplemental Figure 1. Validation of CD8 Subsets from Lung Cancer Patients.** Single cell RNA sequencing reanalysis from Guo et al. 2018[1]. Left, original Psuedotime analysis of 6 CD8$^{+}$ T cell clusters identified by Guo and colleagues, Right, reanalysis of the data set.

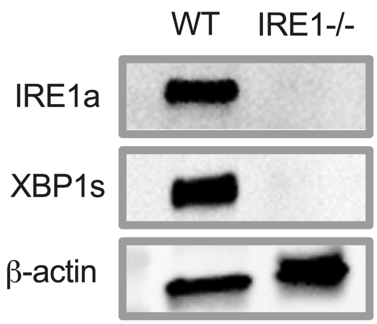


**Supplemental Figure 2**. **Confirmation of IRE1 knockout.** Western blot of 7 day expanded OT-1 T cells receiving electroporation with IRE1 guide RNA (right) or non-targeted control (left) on day 3 post activation.

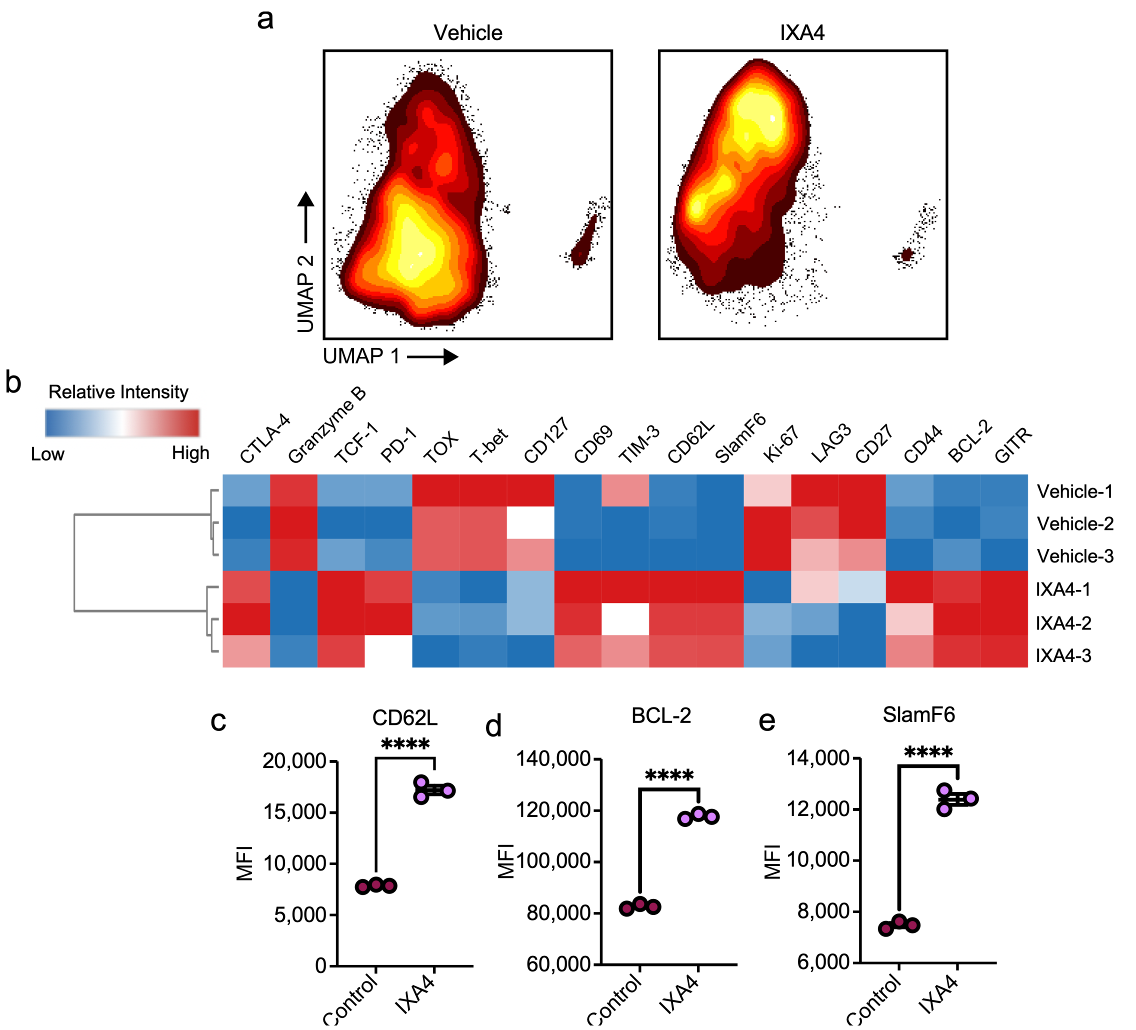


**Supplemental Figure 3. IXA4 induced phenotypic changes.** Flow cytometric analysis of 7-day expanded OT-1 T cells treated with IXA4 (30nM) or vehicle control. a), visualization of differences using the UMAP dimension reduction. b) clustered heatmap of median marker expression per sample. c-e) graphical representation of MFIs from select markers displayed in b. c-e) data are presented as mean ± SEM. Significance determined from unpaired student's *t* test, **** $p < 0.0001$

1. Guo X, Zhang Y, Zheng L, Zheng C, Song J, Zhang Q, Kang B, Liu Z, Jin L, Xing R, Gao R, Zhang L, Dong M, Hu X, Ren X, Kirchhoff D, Roider HG, Yan T, Zhang Z. Global characterization of T cells in non-small-cell lung cancer by single-cell sequencing. Nat Med. 2018;24(7):978-85. Epub 20180625. doi: 10.1038/s41591-018-0045-3. PubMed PMID: 29942094.
